# The role of pro- and antiangiogenic factors in angiogenesis process by Raman spectroscopy

**DOI:** 10.1101/2021.07.02.450827

**Authors:** M. Kopec, H. Abramczyk

## Abstract

Raman spectroscopy and Raman imaging are powerful techniques to monitor biochemical composition around blood vessel. The aim of this study was to understand the role of pro- and antiangiogenic factors in angiogenesis process. Raman imaging and Raman single spectrum measurements allow the diagnosis of cancer biochemical changes in blood vessel based on several biomarkers simultaneously. We have demonstrated that Raman imaging combined with statistical methods are useful to monitoring pro- and antiangiogenic factors responsible for angiogenesis process. In this paper Raman markers of proangiogenic and antiangiogenic factors were identified based on their vibrational signatures. Obtained results can help understand how growing tumor create its vascular system.

**Graphical Abstract:** 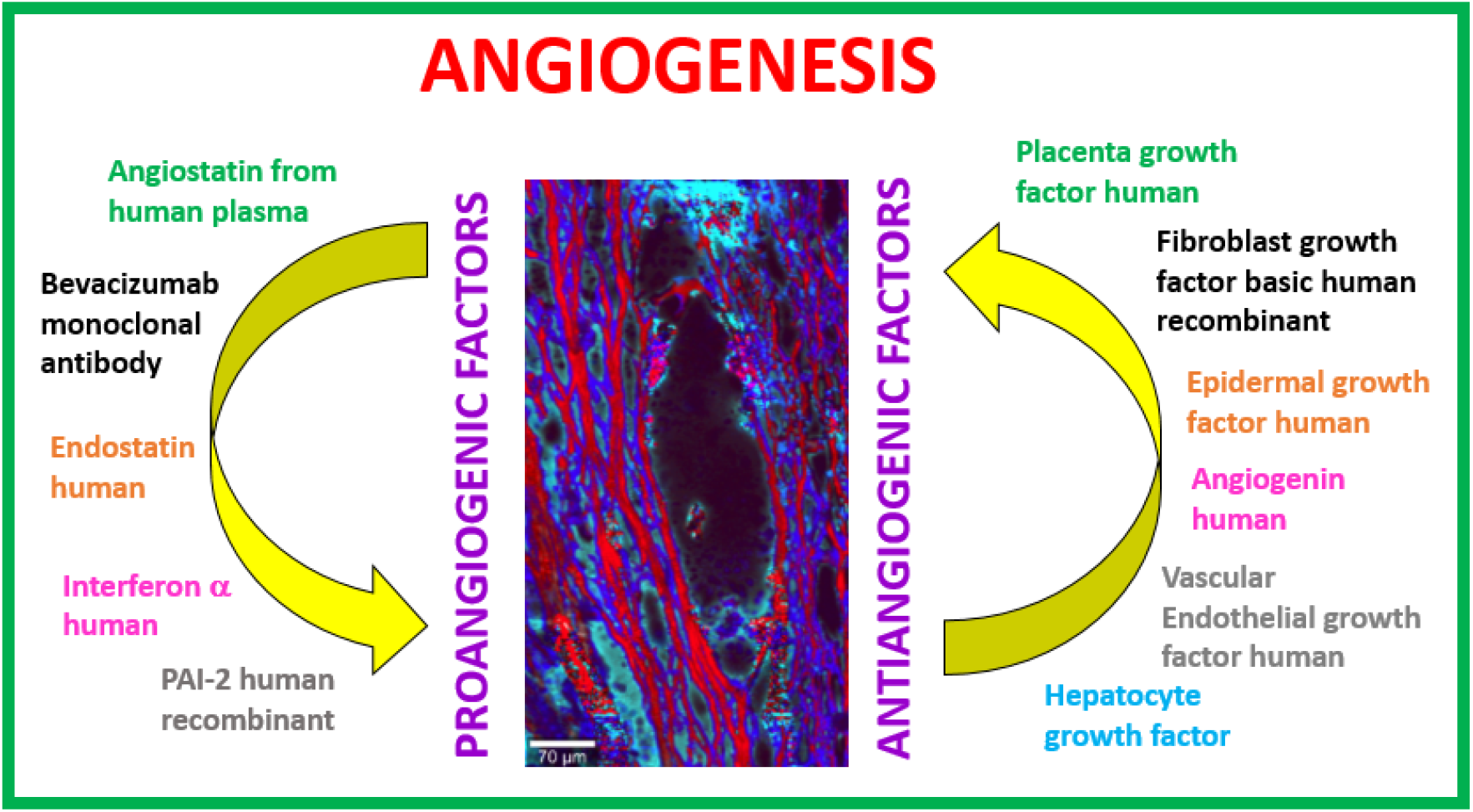

## 1. Introduction

The process of formation new blood vessels is multi-stage. ^1 2 3^ In the first stage the stable vessels (Figure 1A) undergo a vascular permeability increase. This process allows extravasation of plasma proteins (Figure 1B). In subsequent stage the basement membrane is degraded by matrix-metalloproteases and the pericyte-endothelial cell bridging are relived. Due to this fact extracellular matrix liberate sequestered growth factors (Figure 1C). In the next stage endothelial cells proliferate and migrate to their final destination (Figure 1D) and assemble with each other as lumen-bearing cords (Figure 1E). ^4^

**Fig 1.**
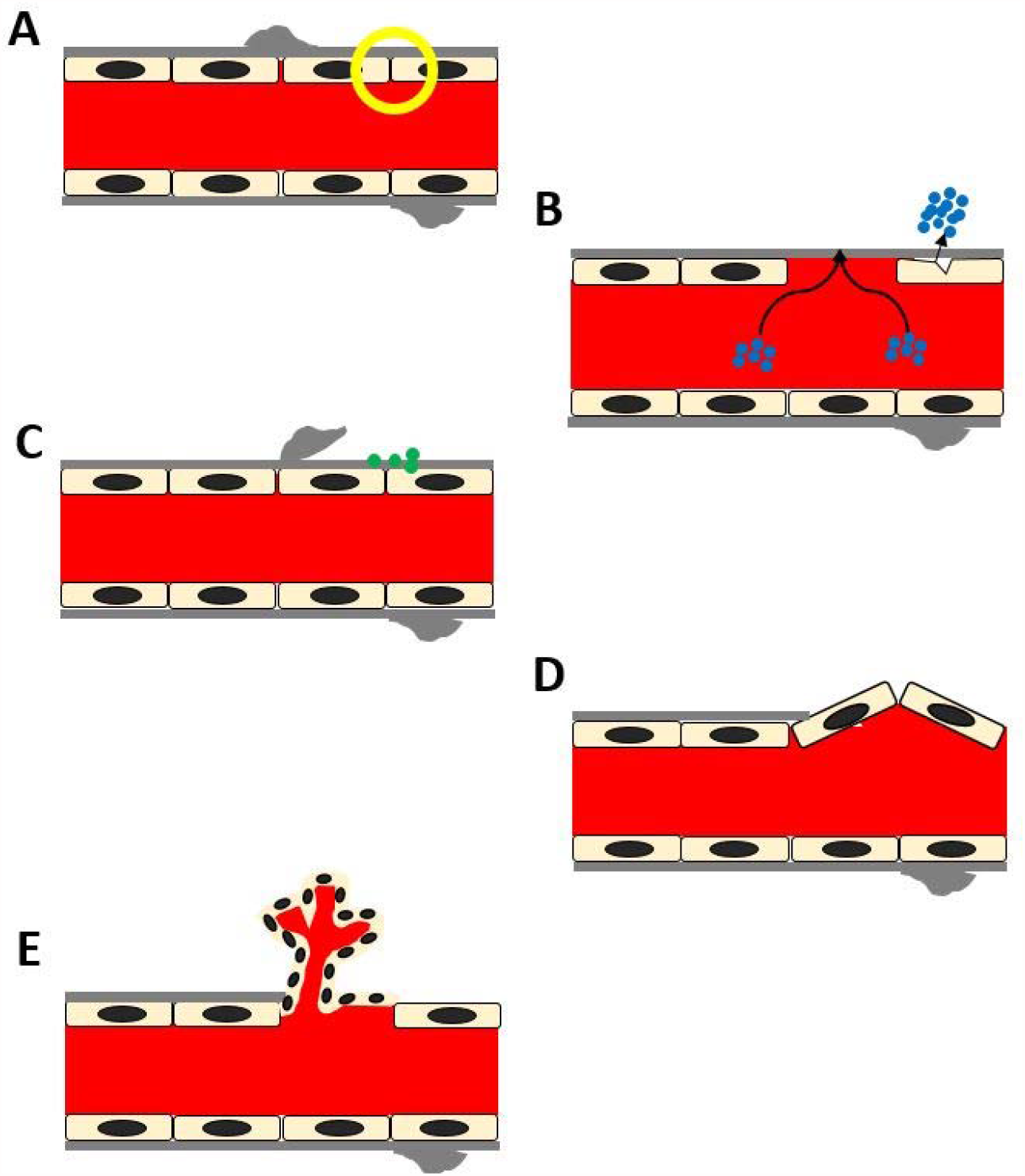
The process of angiogenesis

Figure 1 presents the angiogenesis process.

Every step of angiogenesis process are largely controlled by the proangiogenic and antiangiogenic factors. The role of activators and inhibitors of angiogenesis process is described in numerous papers. ^5 6 7 8 9 10 11 12 13^

Table 1 presents the proangiogenic and antiangiogenic factors in angiogenesis phase.

**Table 1.** Activators and inhibitors of angiogenesis process

| phase | Proangiogenic factor | Antiangiogenic factor |
| --- | --- | --- |
| Degradation of basement membrane | urokinase plasminogen activator (uPA), tissue plasminogen activator (tPA), metalloproteinases (MMPs) | tissue inhibitors of metalloproteinases (TIMPs), Plasminogen Activator Inhibitor (PAI) |
| Migration of endothelial cells | vascular endothelial growth factor (VEGF-A, VEGF-B, VEGF-C, VEGF-D) | Thrombospondin, angiostatin |
| Proliferation of endothelial cells | platelet-derived growth factor (PDGF), platelet-derived endothelial cell growth factor (PDEC GF), fibroblast growth factor (FGF) | Endostatin, prolactin |
| Creating lumen-bearing cords | Angiopoietin 1, tumor growth factor (TGF-α), epidermal growth factor (EGF), angiogenin | Interferons, angiopoietin 2 |

Formation of new blood vessels is regulated by activity of proangiogenic and antiangiogenic factors. Understanding the mechanism of activators and inhibitors functioning in angiogenesis process is very important. ^14 15^ Deregulation of the balance between above-mentioned factors causes pathologic conditions such as cancer development. ^16 17^ This hypothesis for the first time was described by Folkman.^18^ Growing tumor cannot live without oxygen and nutrient.

Tumor use vessels to grow and to delivery products, which are necessary to live and to be up and running.

## 2. Results and discussion

The aim of this paper understands the role of proangiogenic and antiangiogenic factors in angiogenesis process. In this study we analyzed the biochemical composition around blood vessel. To monitor the biochemical changes we used Raman spectroscopy and Raman imaging.

Firstly we focused on analysis of the cross-section of blood vessel. To come to know the biochemical composition we used Raman spectroscopy and Raman imaging. Figure 1 presents the microscopy image, Raman image and Raman spectra in fingerprint and in high frequency regions of the blood vessel.

One can see from Figure 2 (A,B,D) that between obtained Raman images and microscopy image is a perfect match. Raman spectroscopy affords not only perfect reproducing of morphological features but also biochemical characterization of the sample. In our previous publication ^19^ we presented detailed biochemical information about cross-section through vessels. Briefly, in Figure 2 on panel C and E we have shown the characteristic Raman spectra for lipids (blue color), protein (red color) and glycans (turquoise color). ^19^

**Fig 2.**
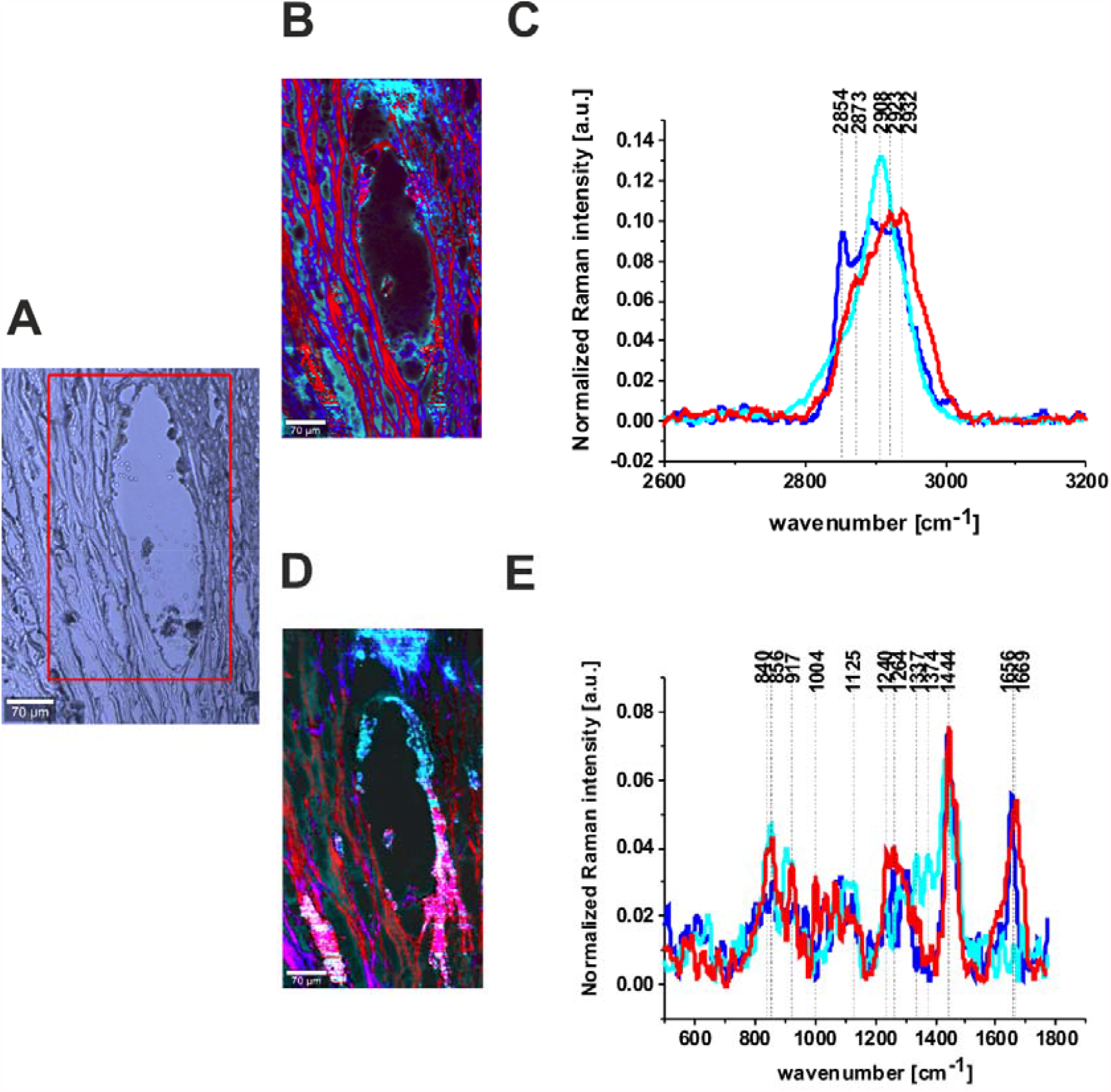
Stitching microscopy image (A), Raman images (360 μm × 660 μm) (B), as well as the normalized (model: divided by norm) Raman spectra (C) in the high frequency region and Raman images (D), normalized (model: divided by norm) Raman spectra (E) in the fingerprint region of the cancerous breast tissue (P149), integration time 0.3 s in high frequency region and 0.5 in the fingerprint region, resolution: 0.5 μm, laser excitation power: 10 mW. Objective 40×, 6 μm-thick slices on calcium fluoride window (CaF2, 25 × 1 mm). The line colors of the spectra correspond to the colors of the Raman maps.

In this paper we want to extend this analysis to explain the role of proangiogenic and antiangiogenic factors in angiogenesis process. To achieve this goal we have registered Raman spectra of pure proangiogenic and antiangiogenic factors.

In Figure 3 we present the Raman spectra of pure proangiogenic factors: angiostatin from human plasma, bevacizumab Monoclonal Antibody, endostatin human; Interferon-a human; PAI-2 human Plasminogen Activator Inhibitor-2.

**Fig 3.**
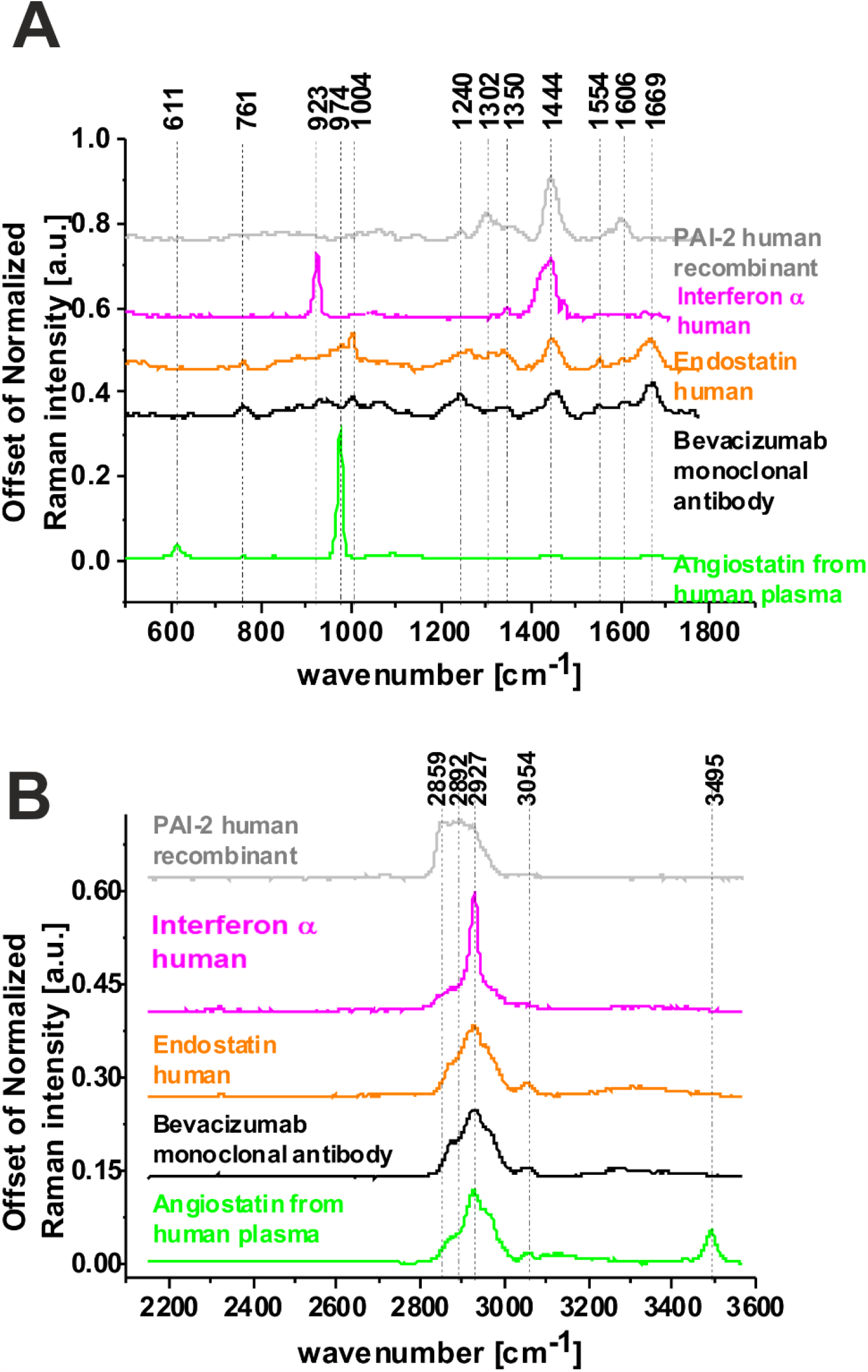
Raman spectra of proangiogenic factors: angiostatin from human plasma (green line), bevacizumab Monoclonal Antibody (black line), endostatin human (orange line); Interferon-a human (magenta line); PAI-2 human Plasminogen Activator Inhibitor-2 (gray line); in fingerprint region (A) and in high frequency region (B)

We prepared the same analysis for antiangiogenic factors. Figure 4 presents Raman spectra of pure antiangiogenic factors: Placenta Growth Factor human, Fibroblast Growth Factor basic human recombinant animal-free, hEGF Epidermal Growth Factor Human, Angiogenin human, Vascular Endothelial Growth Factor human, Hepatocyte Growth Factor Human, which play the role in the angiogenesis process.

**Fig 4.**
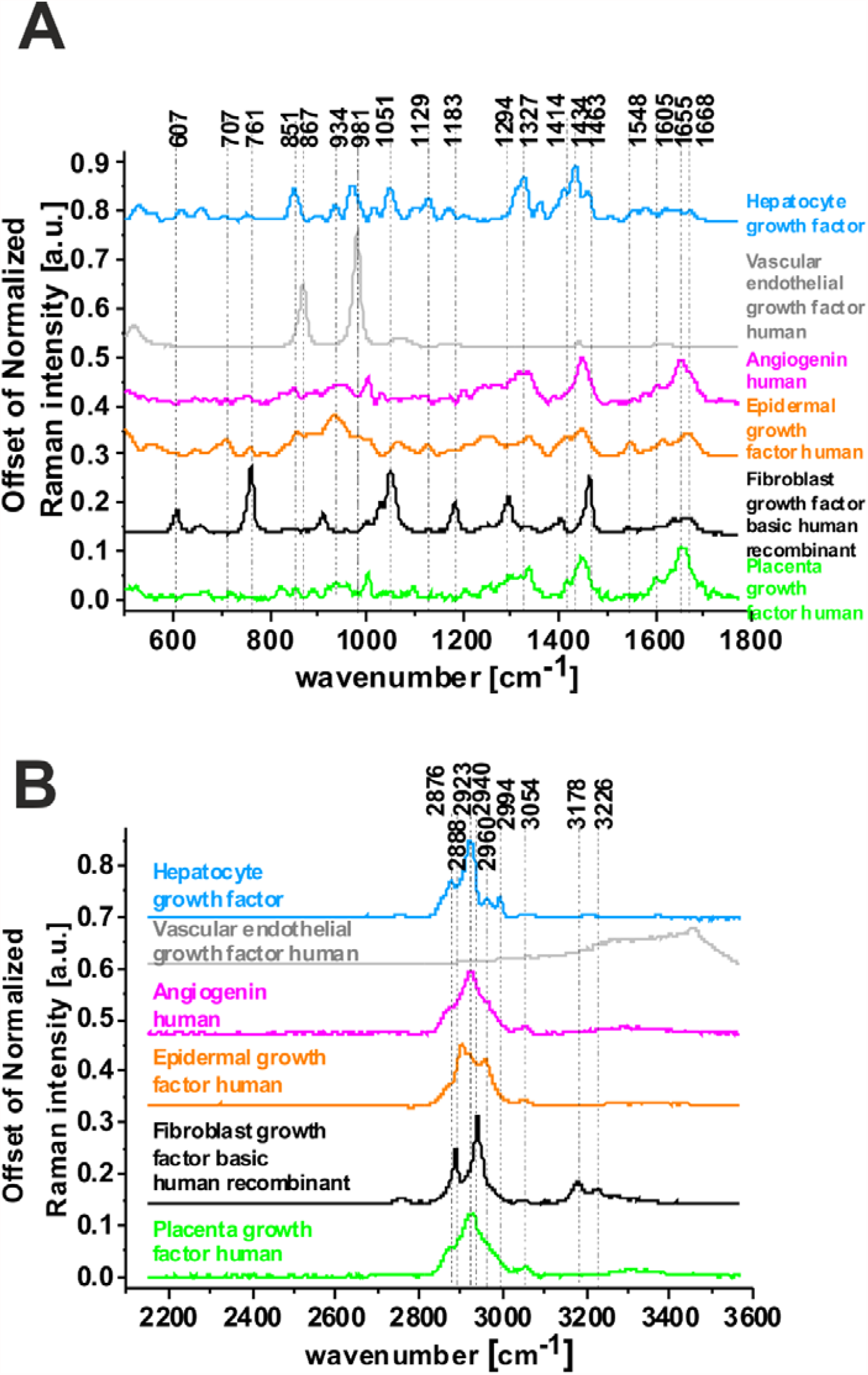
Raman spectra of antiangiogenic factors: Placenta Growth Factor human (green line), Fibroblast Growth Factor basic human recombinant animal-free (black line), hEGF Epidermal Growth Factor Human (orange line), Angiogenin human (magenta line), Vascular Endothelial Growth Factor human (gray line), Hepatocyte Growth Factor Human (light blue line) in fingerprint region (A) and in high frequency region (B)

To identify the proangiogenic and antiangiogenic factors corresponding to the Raman profile in blood vessel we have compared the average Raman spectrum of blood vessel with the pure Raman spectra of proangiogenic factors (angiostatin from human plasma, bevacizumab Monoclonal Antibody, endostatin human, Interferon-a human, PAI-2 human Plasminogen Activator Inhibitor-2) and antiangiogenic factors (Placenta Growth Factor human, Fibroblast Growth Factor basic human recombinant animal-free), hEGF Epidermal Growth Factor Human, Angiogenin human, Vascular Endothelial Growth Factor human and Hepatocyte Growth Factor Human). These results are presented in Figures 5 and 6.

**Fig 5.**
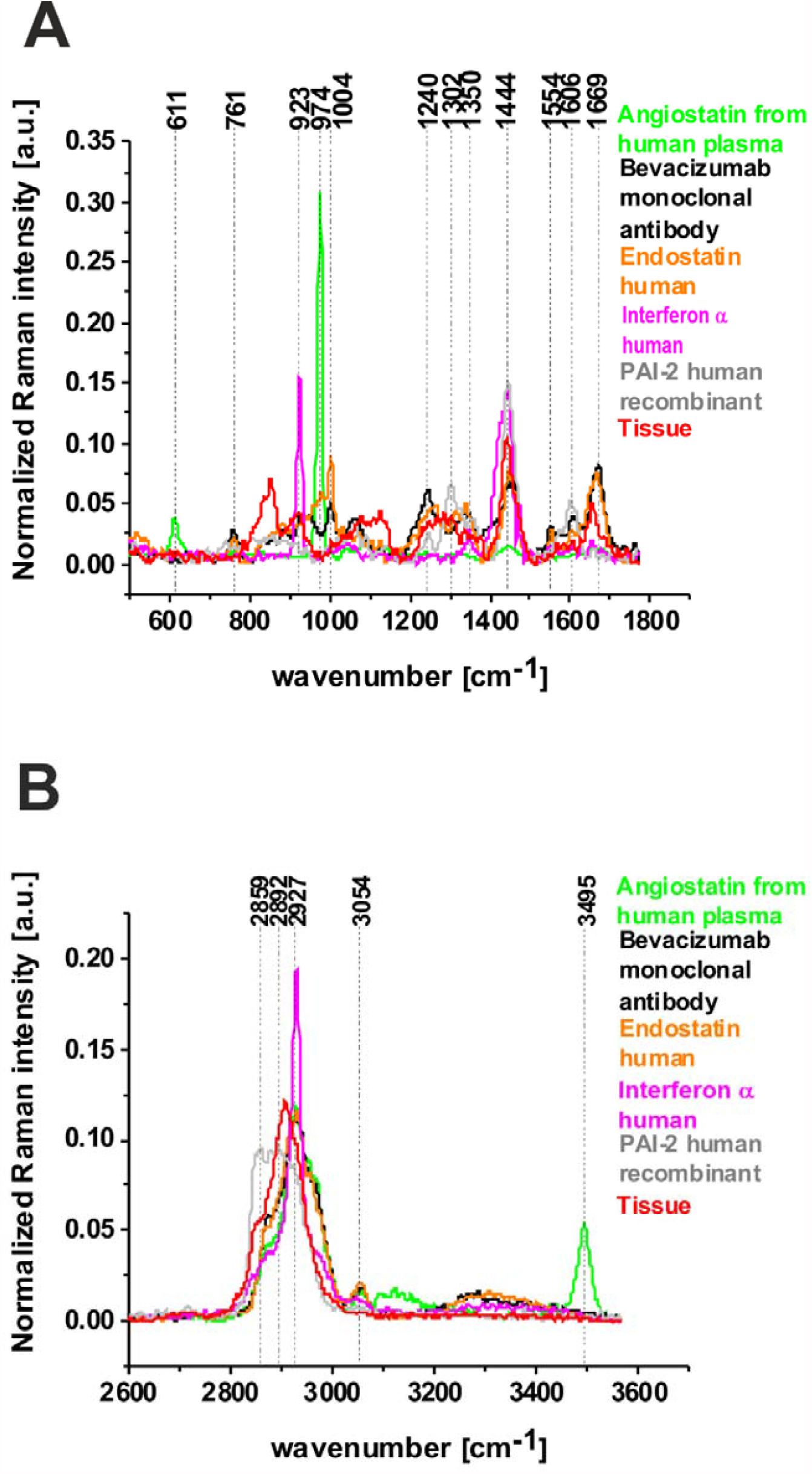
The comparison of average Raman spectra from blood vessels (red line) with the Raman spectra of proangiogenic factors: angiostatin from human plasma (green line), bevacizumab Monoclonal Antibody (black line), endostatin human (orange line); Interferon-a human (magenta line); PAI-2 human Plasminogen Activator Inhibitor-2 (gray line) in fingerprint region (A) and in high frequency region (B).

**Fig 6.**
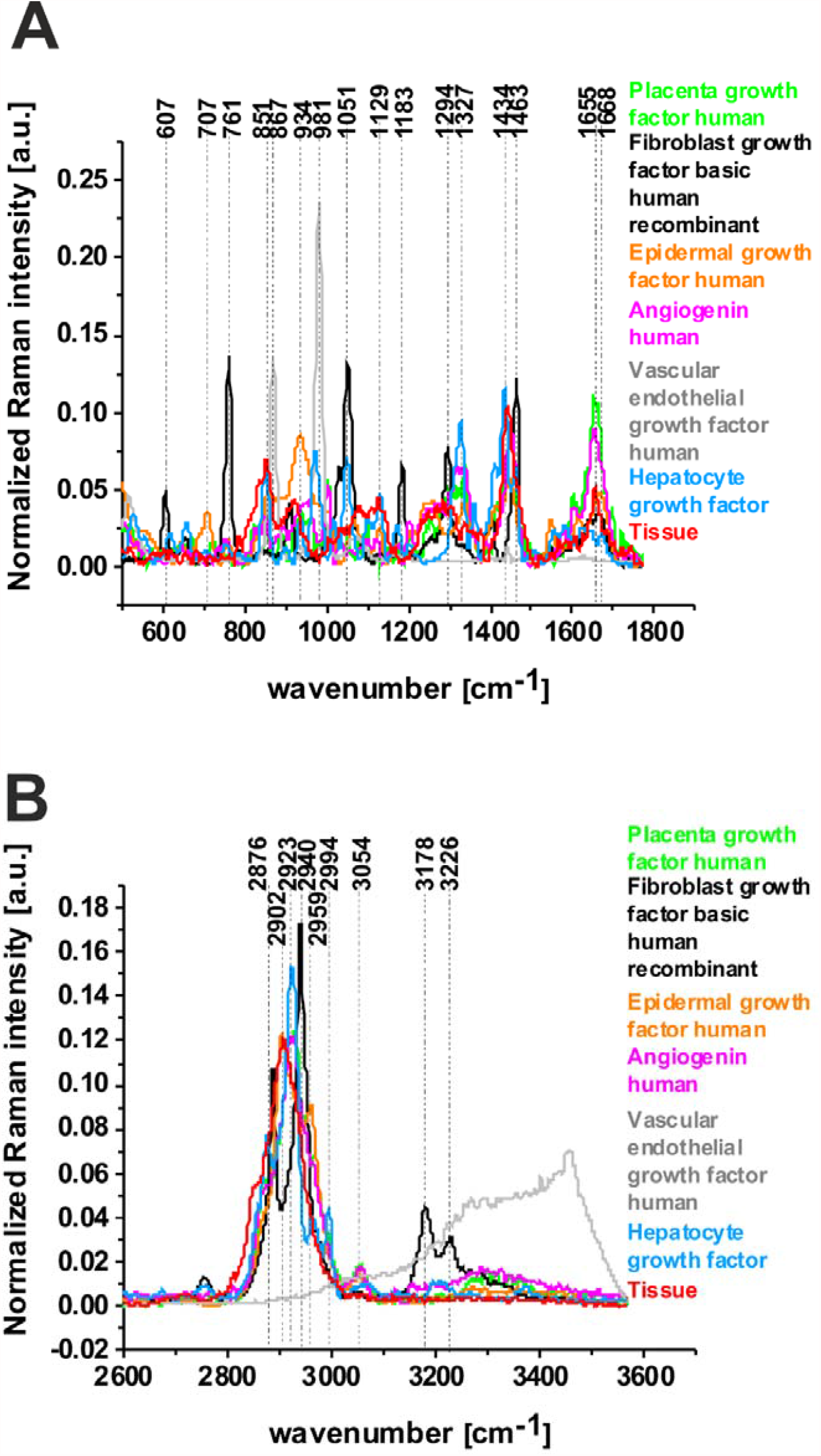
The comparison of average Raman spectra from blood vessels (red line) with the Raman spectra of antiangiogenic factors: Placenta Growth Factor human (green line), Fibroblast Growth Factor basic human recombinant animal-free (black line), hEGF Epidermal Growth Factor Human (orange line), Angiogenin human (magenta line), Vascular Endothelial Growth Factor human (gray line), Hepatocyte Growth Factor Human (light blue line) in fingerprint region (A) and in high frequency region (B).

Figure 5 present comparison between average Raman spectra of blood vessels with Raman spectra of proangiogenic factors.

Extended analysis of Figures 5 shows that the best reproduction of the vessel is obtained for bevacizumab Monoclonal Antibody, endostatin human, Interferon-a human and PAI-2 in fingerprint region. The vibrations present in human tissue at 1240, 1264, 1444, 1669, 2854, 2923 cm^-1^ are assigned to GAG S=O stretching; fatty acids, =C-H bend; fatty acids, triglycerides, CH_2_ or CH_3_ deformations; proteins amide I turn/unsaturated fatty acids, (C=O) stretching, (C-H) def./(C=C) str., collagen, elastin; fatty acids, triglycerides, CH_2_ symmetric stretching and for proteins, respectively. ^20 21 22 23^

To better understanding of the biochemical distribution antiangiogenic factors in blood vessel we prepared the same analysis. Figure 6 present comparison between average Raman spectra of blood vessels with Raman spectra of antiangiogenic factors.

One can see from Figure 6 that the best spectral assignment between Raman spectra from blood vessel and Raman spectra from pure antiangiogenic factors is visible for: Placenta Growth Factor human, hEGF Epidermal Growth Factor Human, Angiogenin human and Hepatocyte Growth Factor Human in high frequency region and for hEGF and Angiogenin human for high frequency region.

To check the similarities between Raman spectrum of human blood vessel and Raman spectra of pure proangiogenic and antiangiogenic factors the correlation analysis was performed. In the Pearson correlation analysis, we took into account the average Raman spectrum for tissue and Raman spectra for pure proangiogenic and antiangiogenic factors, so the presented Pearson correlation coefficient corresponds to the comparison to the whole Raman spectra. This results are presented in Table 2.

**Table 2.** The Pearson correlation coefficients obtained for comparison of the average Raman spectra from blood vessels and the Raman spectra characteristic for pure proangiogenic factors: angiostatin from human plasma, bevacizumab Monoclonal Antibody, endostatin human, Interferon-a human, PAI-2 human Plasminogen Activator Inhibitor-2 and antiangiogenic factors: Placenta Growth Factor human, Fibroblast Growth Factor basic human recombinant animal-free, hEGF Epidermal Growth Factor Human, Angiogenin human, Vascular Endothelial Growth Factor human and Hepatocyte Growth Factor Human

| proangiogenic factors |  |  | antiangiogenic factors |  |  |
| --- | --- | --- | --- | --- | --- |
|  | Pearson correlation coefficient | p-value |  | Pearson correlation coefficient | p-value |
| angiostatin from human plasma | 0.416 | <0.05 | Placenta Growth Factor human | 0.789 | <0.05 |
| bevacizumab Monoclonal Antibody | 0.796 | <0.05 | Fibroblast Growth Factor basic human recombinant animal-free | 0.553 | <0.05 |
| endostatin human | 0.791 | <0.05 | hEGF Epidermal Growth Factor Human | 0.784 | <0.05 |
| Interferon-α human | 0.717 | <0.05 | Angiogenin human | 0.814 | <0.05 |
| PAI-2 human Plasminogen Activator Inhibitor-2 | 0.863 | <0.05 | Vascular Endothelial Growth Factor human | 0.124 | <0.05 |
|  |  |  | Hepatocyte Growth Factor Human | 0.768 | <0.05 |

From Table 2 one can see that the best match between is for PAI-2 (Pearson correlation coefficient was equal 0.863 at the confidence level 0.95), bevacizumab Monoclonal Antibody (Pearson correlation coefficient was equal 0.796 at the confidence level 0.95) and for endostatin human (Pearson correlation coefficient was equal 0.791 at the confidence level 0.95). In contrast to the proangiogenic factors, antiangiogenic factors show that the best match between the average Raman spectra from blood vessels and the Raman spectra characteristic for pure antiangiogenic factors is for Angiogenin human (Pearson correlation coefficient was equal 0.814 at the confidence level 0.95), placenta Growth Factor human (Pearson correlation coefficient was equal 0.789 at the confidence level 0.95) and for hEGF Epidermal Growth Factor Human (Pearson correlation coefficient was equal 0.784 at the confidence level 0.95).

In order to graphical visualize chemical similarities and differences between proangiogenic, antiangiogenic factors and Raman spectrum of blood vessel we have using multivariate statistical methods for data interpretation. Fig. 7 shows the PCA and PLSDA score plot for the Raman spectra of proangiogenic, antiangiogenic factors and Raman spectrum of blood vessel. The values of sensitivity and specificity obtained directly from PLSDA are presented in Table 3.

**Table 3.** The value of sensitivity and specificity for calibration and cross validation procedure from PLS-DA analysis.

| **proangiogenic factors** |  |  |  |  |  |
| --- | --- | --- | --- | --- | --- |
|  | **angiostatin from**<br>**human plasma** | **bevacizumab**<br>**monoclonal antibody** | **endostatin human** | **interferon-α human** | **PAI-2 human**<br>**plasminogen**<br>**activator inhibitor-2** |
| Sensitivity (calibration) | 1.00 | 1.00 | 1.00 | 1.00 | 1.00 |
| Specificity (calibration) | 0.909 | 0.909 | 0.818 | 1.00 | 0.909 |
| Sensitivity (cross validation) | 1.00 | 1.00 | 1.00 | 1.00 | 1.00 |
| Specificity (cross validation) | 0.909 | 1.00 | 1.00 | 1.00 | 1.00 |
| **antiangiogenic factors** |  |  |  |  |  |

|  | placenta growth factor human | fibroblast growth factor basic human recombinant animal-free | hEGF epidermal growth factor human | angiogenin human | vascular endothelial growth factor human | hepatocyte growth factor human |
| --- | --- | --- | --- | --- | --- | --- |
| Sensitivity (calibration) | 1.00 | 1.00 | 1.00 | 1.00 | 1.00 | 1.00 |
| Specificity (calibration) | 0.818 | 1.00 | 0.091 | 0.727 | 1.00 | 0.818 |
| Sensitivity (cross validation) | 1.00 | 1.00 | 1.00 | 1.00 | 1.00 | 1.00 |
| Specificity (cross validation) | 0.909 | 1.00 | 0.636 | 0.909 | 0.909 | 1.00 |

**Fig 7.**
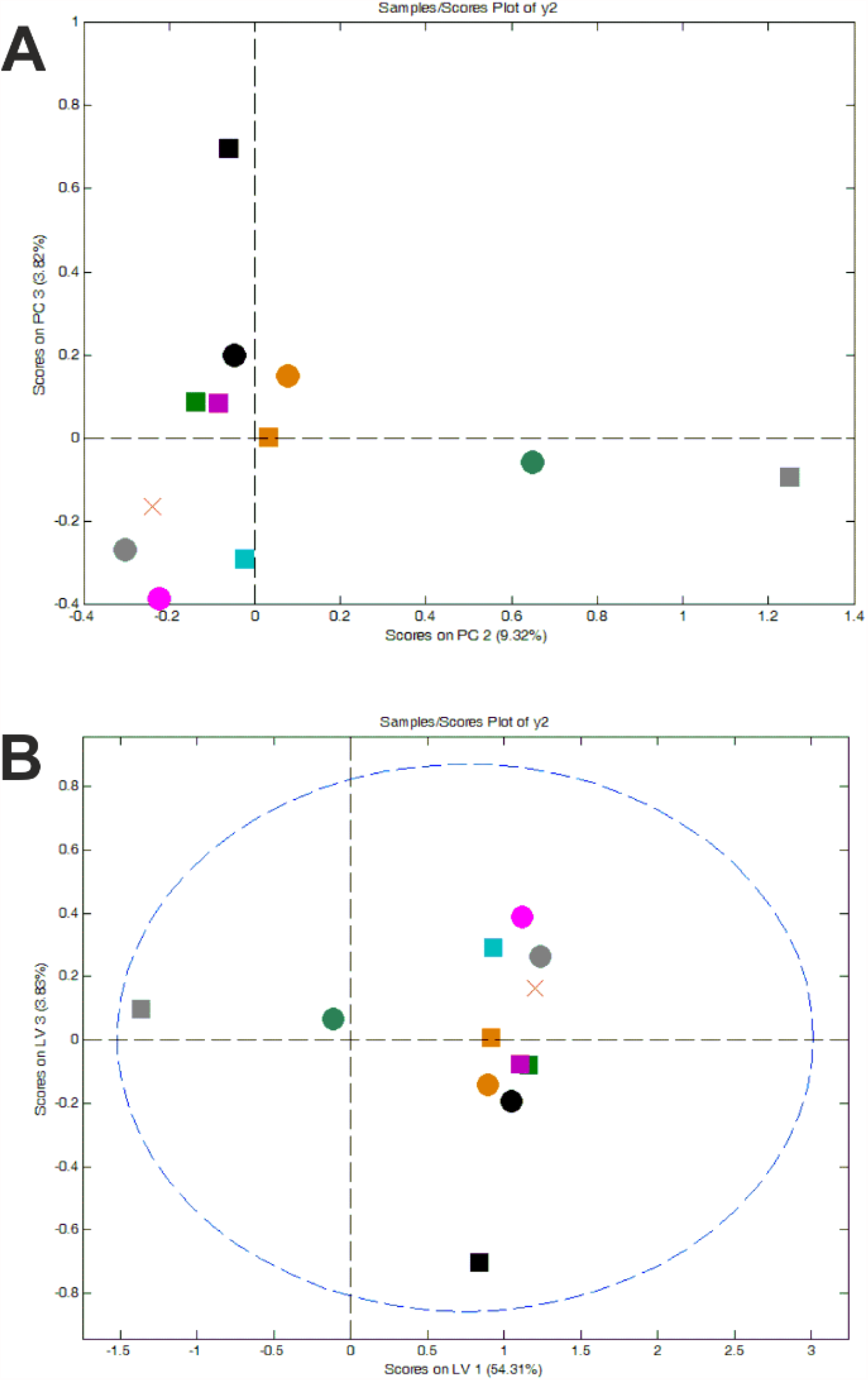
PCA and PLSDA score plot for the Raman spectra of proangiogenic factors: angiostatin from human plasma (green circle), bevacizumab monoclonal antibody (black circle), endostatin human (orange circle), Interferon-a human (magenta circle), PAI-2 human plasminogen activator inhibitor-2 (gray circle); antiangiogenic factors: Placenta Growth Factor human (green square), Fibroblast Growth Factor basic human recombinant animal-free (black square), hEGF Epidermal Growth Factor Human (orange square), Angiogenin human (magenta square), Vascular Endothelial Growth Factor human (gray square) and Hepatocyte Growth Factor Human (turquoise square); average Raman spectra from blood vessels (red cross)

By plotting the principal component scores, similarities between the samples can be revealed. One can see from Fig 7 that among proangiogenic factors angiostatin from human plasma (green circle) is the most separated from Raman spectrum of blood vessel (red cross). The Pearson correlation coefficient is only 0.416. While among antiangiogenic factors Vascular Endothelial Growth Factor human (gray square) and Fibroblast Growth Factor basic human recombinant animal-free (black square) are the most separated from Raman spectrum of blood vessel. The Pearson correlation coefficient is only 0.124 and 0.553 respectively.

Moreover, Raman spectrum for Interferon-a human (magenta circle), PAI-2 human plasminogen activator inhibitor-2 (gray circle) and for Hepatocyte Growth Factor Human (turquoise square) are the most grouped with Raman spectrum for blood vessel. This result was presented also in Table 2. Correlation analysis shows the most perfect match between Raman spectrum of blood vessel and in Raman spectra of pure factors (Pearson correlation coefficient is 0.717, 0.863 and 0.768 respectively at the confidence level 0.95).

Table 3 present the values for sensitivity and specificity obtained from chemometric methods. The high values for sensitivity and specificity highlight the importance of Raman spectroscopy and imaging as a new diagnostic tool in angiogenesis process.

## Conclusions

We have demonstrated that Raman spectroscopy and Raman imaging allows to study the biochemical composition of human blood vessels. Profile characteristic for proteins, glycans and lipids have been mapped in human breast tissue. This paper examined pure proangiogenic and antiangiogenic factors by Raman spectroscopy. Based on Raman spectra for proangiogenic and antiangiogenic factors we identified the factors, which are responsible for angiogenesis process. Results presented in this manuscript suggest that among antiangiogenic factors Placenta Growth Factor human, hEGF Epidermal Growth Factor Human, Angiogenin human and Hepatocyte Growth Factor Human are the candidates for Raman biomarkers in angiogenesis process. The proangiogenic factors which take part in angiogenesis process were also identified, it was: bevacizumab Monoclonal Antibody, endostatin human, Interferon-α human and PAI-2. The results presented in the paper demonstrate that Raman spectroscopy and Raman imaging are powerful methods that will bring an important contribution to the understanding the role of proangiogenic and antiangiogenic factors in angiogenesis process.

## Material and methods

### Chemicals

All chemicals compounds have been purchased from Sigma-Aldrich: angiostatin from human plasma (SRP6044; Sigma Aldrich), bevacizumab Monoclonal Antibody (MSQC20; Sigma

Aldrich), endostatin human (SRP3031, Sigma Aldrich); Interferon-a human (I4276; Sigma Aldrich); PAI-2 human Plasminogen Activator Inhibitor-2 (SRP3137; Sigma Aldrich); Placenta Growth Factor human (P1588; Sigma Aldrich); Fibroblast Growth Factor basic human recombinant animal-free (GF003AF; Sigma Aldrich); hEGF Epidermal Growth Factor Human (E9644; Sigma Aldrich); Angiogenin human (A6955; Sigma Aldrich); Vascular Endothelial Growth Factor human (V7259; Sigma Aldrich); Hepatocyte Growth Factor Human (H1404; Sigma Aldrich)

### Tissue preparation

The tissues were collected during operation at the WWCOiT Nicolaus Copernicus in Lodz. Fresh, non-fixed samples were cut into 6 um slices and put on CaF_2_ windows. All the experiments were approved by the institutional Bioethical Committee at the Medical University of Lodz, Poland (RNN/323/17/KE/17/10/2017). Written consents from patients or from legal guardians of patients were obtained. Spectroscopic analysis did not affect the scope of surgery and course and type of undertaken hospital treatment.

### Raman spectroscopy and Raman imaging

All Raman results presented in this manuscript were performed using WITec alpha 300 RSA+ combined with confocal microscope coupled via the fibre of a 50 mm core diameter with an UHTS (Ultra High Throughput Spectrometer) spectrometer and a CCD Camera (Andor Newton DU970N-UVB-353) operating in standard mode with 1600×200 pixels at 60C with full vertical binning. To obtain all presented results we used laser diode (SHG of the Nd:YAG laser (532 nm)) and 40x dry objective (Nikon, objective type CFI Plan Fluor CELWD DIC-M, numerical aperture (NA) of 0.60 and a 3.6–2.8 mm working distance). All Raman experiment were performed using laser with a power 10 mW at the sample position. To acquisition and preprocess of the data (cosmic rays removing, smoothing and removing background) we used WITec Project Plus. Detailed methodology is available elsewhere ^24 25 26^

### Statistical Analysis

To show the match between the average Raman spectra from blood vessels and the Raman spectra of pure proangiogenic factors and antiangiogenic factors correlation analysis (Pearson correlation coefficient at the confidence level 0.95) was performed in Origin.

Principal Components Analysis and Partial Least Squares Discriminant Analysis was performed using Mathlab and PLS_Toolbox Version 4.0. Detailed description of chemometric methods was presented in our previous paper. ^27 28^

## Acknowledgements

This work was supported by the National Science Center of Poland Miniatura 4 (grant 2020/04/X/ST4/00325) and UMO-2019/33/B/ST4/01961.

## Author Contributions

Conceptualization: M.K.; Funding acquisition: M.K., H.A., Investigation: M.K. Methodology: M.K., H.A.: Writing-original draft: M.K.; Manuscript editing: M.K., H.A. All authors have read and agreed to the published version of the manuscript.

